# Intrauterine growth restriction causes cellular, molecular, and behavioral deficits consistent with abnormal dentate gyrus neurogenesis in mice

**DOI:** 10.1101/2020.07.16.207449

**Authors:** Ashley S. Brown, Matthew Wieben, Shelby Murdock, Jill Chang, Maria Dizon, Richard I. Dorsky, Camille M. Fung

## Abstract

**Background:** Children born with intrauterine growth restriction (IUGR) are at increased risk for cognitive impairment including learning and memory deficits. Dentate gyrus (DG) granule neurons relay cortical information into the hippocampus proper for memory formation, and their production is highly dependent on environmental signals. However, it is unknown whether IUGR affects DG neurogenesis, and thus provides a potential mechanism underlying abnormal learning and memory function.

**Methods:** Using a hypertensive disease of pregnancy mouse model of IUGR, we assessed multiple behaviors, quantified neural stem and progenitor cells (NSPCs) and developing neurons in the DG, and characterized transcriptional effects on molecular pathways in the hippocampus.

**Results:** We found that the predominant behavioral phenotype in IUGR offspring, short-term implicit learning and memory deficits, was associated with accelerated DG neurogenesis and NSPC depletion. Consistent with known molecular regulators of DG neurogenesis, we also found strong evidence for decreased Wnt pathway activity following IUGR.

**Conclusion:** We have discovered that postnatal memory deficits are associated with accelerated NSPC differentiation following IUGR, a phenotype that could be explained by decreased Wnt signaling.

## Introduction

Intrauterine growth restriction (IUGR) describes a condition of fetal weight below the genetically predetermined potential (1). In high-income countries, IUGR complicates 3-9% of all pregnancies whereas the incidence is reportedly six-fold higher in low-income countries (2). The etiologies of IUGR are numerous, and may relate to maternal, placental and cord abnormalities, or fetal factors. The primary cause in developed countries is hypertensive disease of pregnancy which creates uteroplacental insufficiency, in which the placenta fails to support fetal growth. IUGR infants are prone to a range of health problems including impairment in motor, sensory, cognitive, and learning and memory tasks (3).

The hippocampus, a medial cortical structure required for forming new memories and learned behaviors (4), is divided into the dentate gyrus (DG) and cornu ammonis (CA) fields 1-3 in both humans and rodents, and is particularly susceptible to IUGR (5). Both IUGR children and animal models of IUGR have shown reductions in hippocampal volume by magnetic resonance imaging and have correlated poorer memory performance with the magnitude of hippocampal volume reduction (6-9). Multiple experimental models of IUGR have also indicated that both the neuron number and dendritic-axonal morphologies within the DG, CA1, and CA3 are altered following uteroplacental insufficiency. For example, guinea pig neonates with IUGR have fewer cells in the CA1 region (8), consistent with reduced cell numbers in CA1 pyramidal layer in juvenile rats with IUGR (9). Furthermore, a reduction in dendritic length and outgrowth, reduced branch numbers of apical dendrites, and alterations in basal dendritic branch points in the CA1 region have been reported (10). What is unknown from existing literature are the *in utero* changes that lead to the functional and cellular differences observed in postnatal life.

We therefore employed our laboratory’s mouse model of IUGR induced by thromboxane A_2_-analog infusion to dissect the cellular and molecular events that are disrupted during embryonic DG neurogenesis (11, 12). We first performed a battery of behavioral assays to establish the functional deficits in this model. We then focused on DG neurogenesis and systematically enumerated the composition of Sox2^+^ neural stem and progenitor cells (NSPCs) and the neuronal production of Tbr2^+^ intermediate neuronal progenitors (INPs), NeuroD^+^ neuronal progenitors (NPs), and Prox1^+^ immature and mature granule neurons during early and late IUGR. Finally, we performed RNA-sequencing to determine the differences in gene expression between sham control and IUGR hippocampi that may be associated with the cellular phenotype.

## Methods

### Mouse model of IUGR

All animal procedures were approved by the University of Utah Animal Care Committee. Full details can be found on model inception (11). Briefly, we set up timed matings of wild-type C57BL/6J mice (000664, The Jackson Laboratory, Bar Harbor, ME). At embryonic day (E) 12.5 (term gestation ∼20 days), pregnant dams were anesthetized and micro-osmotic pumps (1007D, 0.5ul/h, Durect Corporation, Cupertino, CA) containing either vehicle (0.5% ethanol = sham control) or 4000ng/μl of U-46619 (a thromboxane A_2_-analog) dissolved in vehicle (16450, Cayman Chemical, Ann Arbor, MI) were implanted retroperitoneally. Pups used for embryonic studies were delivered via Caesarian sections after maternal anesthesia at E15.5 or E19. Pups used for postnatal behavioral testing were born vaginally. All sham control and IUGR pups were cross-fostered to unmanipulated dams to minimize surgery-related complications. We previously established that mouse dams who received U-46619 developed maternal hypertension within 24 h of pump implantation with mean blood pressures being 20% higher than sham-operated dams. Additionally, IUGR offspring exhibited smaller weight gain from E17.5 to E19 and were 15% symmetrically growth restricted at birth compared to sham controls (11, 12).

### Behavioral Assays

#### Novel Object Recognition

We habituated 2-3 month old sham control and IUGR mice of both sexes (n=5/each treatment and sex) to the test arena (a black box measuring 40 cm ⨯ 40 cm equipped with a camera overhead that allowed for the filming of mouse exploration) for 15 min on day 1. The next day, we placed two identical objects in opposite corners of the arena and mice were allowed to explore for 15 min. The duration of time spent exploring either object minus the time spent climbing on the object (which does not constitute exploration) was recorded with a camera and calculated using EthoVision 3.1 (Noldus Information Technology Inc., Leesburg, VA). On the third day, we substituted one of the old objects with a novel object and mice were allowed to explore for 15 min. The duration of time spent with the old and novel objects subtracting climbing time was calculated using EthoVision.

#### Fear Conditioning

Six-month old sham control and IUGR mice of both sexes (n-5/each treatment and sex) were placed into Plexiglass conditioning chambers on day 1 for 5 trials of a 30 second tone at 75dB with a 20 second trace interval, followed by a brief 1 second mild foot shock at 0.7mA. Mice were returned to the same chamber the next day for 5 min and scored for time spent in freezing motion in the absence of tone in *contextual conditioning*. They were then placed into a novel chamber for 5 min and scored for time spent in freezing motion with the presentation of the tone but without a foot shock in *cued test*.

#### Open field test

We recorded spontaneous activities in an open field for 30 min using the same black square as that used in object interaction/recognition in 2-3 months old mice (n-5/each treatment and sex). The lighting across the floor of the arena was even. Mice were placed into the center of the arena as a starting point. Both horizontal (locomotion) and vertical (rearing) activities were recorded by an overhead camera. Behavioral outputs using EthoVision included total duration (seconds (s)), distance traveled (cm), mean velocity (cm/s), and rearing frequency in the center and periphery of the arena. We also measured latency to the periphery (s) from the center.

#### Elevated plus maze

We used an elevated, plus-shaped (+) apparatus with two open and two enclosed arms (Noldus Information Technology Inc) in 2-3 months old mice (n-5/each treatment and sex). The behavioral model is based on a general aversion of rodents to open spaces. Mice were initially placed in the center of the + facing an open arm. Behavioral outputs using EthoVision included frequency and duration (s) in the center, open, or closed arms as well as the latency (s) to enter either into an open or closed arm from the center for a total recording time of 10 min.

#### Prepulse Inhibition (PPI)

The premise of this test is a normal reduction in startle magnitude when an intense startling stimulus (or pulse) is preceded by a weaker pre-stimulus (or prepulse). It evaluates sensorimotor gating which is the ability of a sensory event to suppress a motor response (13). We first acclimated 2-3 months old mice (n-5/each treatment and sex) for 5 min in the SR-LAB startle apparatus (San Diego Instruments, San Diego, CA). During the acclimation period, a constant 70 dB white noise emitted by the digital sound level meter (FLIR Systems, Extech) was presented for the animal to adapt to the environment. The session then proceeded through the presentation of 60 trials: the first 5 trials were 5 pulse-alone trials where 120 dB of white noise was presented for 20msec (i.e. no prepulse), the intermediate 50 trials were divided into 20msec of randomized pulse-alone trials, prepulse-pulse trials of three different prepulse intensities (73 dB, 76 dB, and 82 dB) above the 70 dB background with inter-trial interval varying between 10 and 20 s intended to minimize habituation to startle across trials, and no stimulus trials as a measure of basal motor activity in the cylinder, and concluding with a final block of pulse-alone trials. We recorded the startle reflex which consisted of involuntary contractions of whole-body musculature in response to the acoustic stimulus using a digitized piezoelectric accelerometer (San Diego Instruments). Behavioral outputs calculated were startle reactivity which was the maximal response peak amplitude to the startle pulse, % PPI with the increasing prepulse intensivities, and Tmax which was the latency time to peak response.

#### Forced swim test

We placed 2-3 months old mice (n-5/each treatment and sex) into cylindrical tanks (30 cm tall x 20 cm diameter) constructed of Plexiglas with room temperature water level of 15 cm from the bottom. We recorded for a total of 20 min from the time mice were placed into the water to assess their mobility. Output variables included the frequency and duration to struggle, and frequency, duration, and latency to treading water and floating.

#### Elevated bridge test

We placed 2-3 months old mice (n-5/each treatment and sex) onto a 1 meter long beam resting 50cm above ground. A 60 watt lamp was used to shine light above the start point and served as an aversive stimulus for them to traverse the beam. A nylon hammock was stretched below the beam to cushion any falls. Latency to move and time to cross the entire beam (both in s) were recorded and quantified.

### Brain perfusion and immunohistochemistry

Pregnant mouse dams at E15.5 or E19 were sedated with ketamine (80-100μg/g) and xylazine (7.5-16μg/g). Once adequate anesthesia was achieved by lack of withdrawal on toe pinch, a thoracotomy was performed to expose the heart. A 25G needle was inserted into the left ventricle and the animal was perfused with normal saline followed by perfusion with 4% paraformaldehyde (PFA). A Caesarian section was performed to deliver pups. Pup brains were extracted and post-fixed in additional 4% PFA for 1 h in room temperature (RT). After washing with PBS, embryonic brains were cryoprotected with increasing concentrations of sucrose and embedded in OCT medium. Brains were stored in -80°C until cryosectioning at 12 μm. All brain sections from the beginning of the dorsal to ventral hippocampus were collected in a series of 20 slides such that the 1^st^, 21^st^, and 41^st^ sections were on the first slide while 2^nd^, 22^nd^, 42^nd^ sections were on the second slide, etc.

For immunofluorescent staining of cell types of interest, we hydrated each slide spanning the dorsal hippocampal DG with 1X PBS and proceeded with antigen retrieval with neutral buffer at 37°C x 30 mins for Sox2, 10mM citrate buffer pH 6.0 at 37°C x 30 mins for Tbr2, neutral buffer at 85°C x 10 mins for NeuroD, and 10mM citrate buffer pH 6.0 at 95°C x 5 mins for Prox1 (neutral buffer CTS016, R&D Systems, Minneapolis, MN). Sections were then blocked with 10% or 5% (for Tbr2) normal goat serum at RT for 1 h. We incubated with the following primary antibodies at 4°C overnight: Sox2 (1:300, AB5603, MilliporeSigma, Burlington, MA), Tbr2 (1:500, ab23345, Abcam, Cambridge, MA), NeuroD (1:100, 4373S, Cell Signaling, Danvers, MA), and Prox1 (1:1000, AB5475, MilliporeSigma). After washing, we incubated with 1:1000-1:2000 goat anti-rabbit secondary antibody, Alexa Fluor 594 (A11037, Thermo Fisher Scientific, Waltham, MA) at RT for 1 h. We applied nuclear DAPI counterstain and added Fluoromount-G (0100-01, SouthernBiotech, Birmingham, AL) before all slides were cover slipped for imaging (n-6-7/each treatment and sex).

#### 5-ethynyl-2’deoxyuridine (EdU) labeling

EdU is a nucleoside analog of thymidine and is incorporated into DNA during active DNA synthesis. Detection is based on a click reaction, a copper-catalyzed covalent reaction between an azide which is the Alexa Fluor 488 dye and an alkyne which is the EdU (C10339, Invitrogen Corporation, Carlsbad, CA). EdU is capable of crossing the placenta to be taken up by actively dividing NSPCs in the fetal brain. We injected 100μg/g EdU for 2 h prior to pregnant dam harvest. Pup brains were processed as noted above. We followed the manufacturer’s protocol for EdU detection.

#### Imaging and Quantification of cells

All sections were imaged using a laser scanning confocal microscope (Olympus, Waltham, MA) with a 20x objective lens and the Fluoview FV1000 software. Once images were obtained, we used NIH ImageJ to manually outline the hippocampal DG area. Images were thresholded to obtain the optimal exposure of each image. The number of blue DAPI^+^ nuclei was quantified within the area. The number of Sox2^+^, Tbr2^+^, NeuroD^+^, Prox1^+^, and EdU^+^ cells were counted as a percentage of total DAPI^+^ cells. Data are expressed as percentages of Sox2^+^, Tbr2^+^, NeuroD^+^, Prox1^+^, or EdU^+^ cells within the DG.

### RNA-sequencing

We dissected E15.5 sham control and IUGR hippocampi of both sexes (n=4/each treatment) from whole mouse brains. Total RNA was extracted, followed by verification of RNA quality and quantity by Agilent ScreenTape Assay at the University of Utah High-Throughput Genomics core facility. RNA sequencing library was made using the Illumina TruSeq Stranded Total RNA Sample Prep Kit with Ribo-Zero Gold which allowed for the removal of cytoplasmic and mitochondrial rRNA. Remaining RNA was chemically fragmented and random primed for reverse transcription to construct cDNA libraries. The average insert size of libraries was ∼150 bp with inserts ranging from 100-400 bp. Once the libraries were made and validated, sequencing adapters were added and sequenced as single-reads using the HiSeq 2500 platform.

Reads were aligned to *Mus musculus* genome assembly GRCm38 (mm10, Genome Reference Consortium). Of the 16,873 genes contained in the mouse genome, we determined the differential expression of protein-coding transcripts that had ≥10 but □50,000 counts between sham control and IUGR hippocampi. Using a false discovery rate of 13 (= p≤0.05) and log_2_ ratios ≥ +1 or ≤ −1 (= two-fold increase or decrease in gene expression), 611 protein-coding transcripts were found to be differentially expressed. We entered these differentially-expressed genes into Ingenuity Pathway Analysis (Qiagen Bioinformatics) to determine canonical pathways that were aberrantly expressed in IUGR. Pathways with –log(p-value) above the threshold denote significance of p≤0.05. The ratio in each pathway signifies the number of genes detected in RNA-seq divided by the number of genes known in that pathway.

### Quantitative real-time polymerase chain reaction (qPCR)

Genomic-free total RNA was isolated from a separate cohort of E15.5 sham control and IUGR hippocampi (n=4-5/each treatment) using Nucleospin RNA protocol (Takara Bio USA, Mountain View, CA). Total RNA was quantified with a Nanodrop spectrophotometer ND-1000 (Wilmington, DE). cDNA was synthesized from 4 μg of total RNA using random hexamers of SuperScript III First-Strand Synthesis System (Invitrogen Corporation). Using real-time qPCR as previously described (11), we examined mRNA levels of Wnt signaling genes (Wnt2b, Wnt3a, Rspo3, Fzd7, Dkk1, Wif1, Sp5) to corroborate with RNA-seq data. All primers and probes were purchased as Taqman Assays-on-Demand with assay identification numbers of Mm00437330_m1 (Wnt2b), Mm00437337_m1 (Wnt3a), Mm01188251_m1 (Rspo3), Mm00433409_s1 (Fzd7), Mm00438422_m1 (Dkk1), Mm00442355_m1 (Wif1), and Mm00491634_m1 (Sp5) (Applied Biosystems, Foster City, CA). Each sample was run in quadruplicates. Relative quantitation comparing 2^-ΔCt^ values was used to analyze changes in mRNA expression between sham control and IUGR using GAPDH as a control. We have previously validated that GAPDH was an appropriate control using parallel serial dilutions between sham control and IUGR cDNA and that the amplification efficiencies between the target genes and GAPDH were comparable.

### Statistics

Data are expressed as means ± SEM. Normality of data and equal variance analyses were performed to determine whether parametric tests were appropriate. Student t-test was used to analyze simple effects for 2 groups. Two-way ANOVA was used to examine the effects of sex and treatment (sham control or IUGR) on effects. Statistical significance was set at p□0.05 using STATVIEW software (SAS Institute Inc., Cary, NC).

## Results

### Behavioral assessment of young adult sham control and IUGR mice (2-3 months of age)

#### Short-Term Implicit Memory Testing: Object Interaction/Recognition and Fear Conditioning

We noted no difference in the duration of time spent with the left or right object during training day (n=5/treatment and sex, Figure 1a). When we substituted one of the old objects with a novel object on the next day, IUGR females spent less time exploring the novel object compared to sham females (n=5/treatment and sex, p<0.05, Figure 1b). IUGR males, on the other hand, explored the novel object as frequently as sham males (n=5/treatment and sex, Figure 1b). For fear conditioning, both IUGR females and males showed less freezing time or fear during both contextual conditioning and cued test when compared to sex-matched sham controls (n=5/sex/group, Figures 2a & b).

**Figure 1.**
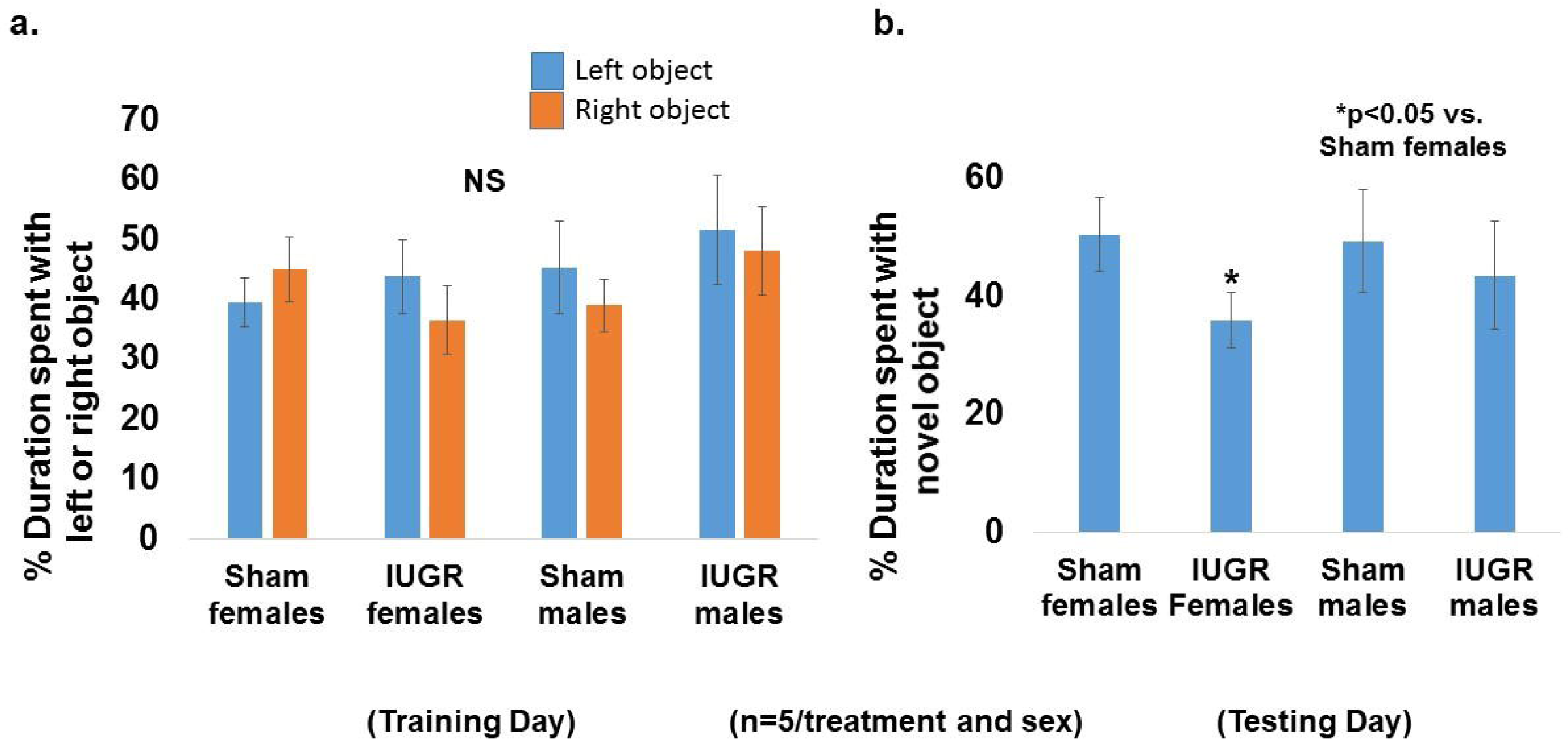
(a) Percentage of duration of time spent with left or right object in training day. Blue bars denote the left object. Orange bars denote the right object. Sham and IUGR offspring spent equal duration of time exploring left and right objects. (b) Percentage of duration of time spent with novel object in testing day. IUGR females spent less time with the novel object, i.e. more time with the old object, demonstrating a lack of memory to the familiar object from previous day’s exposure. *p<0.05 compared to sham females. NS = non-significance. n=5 for each treatment (sham or IUGR) and sex (male or female).

**Figure 2.**
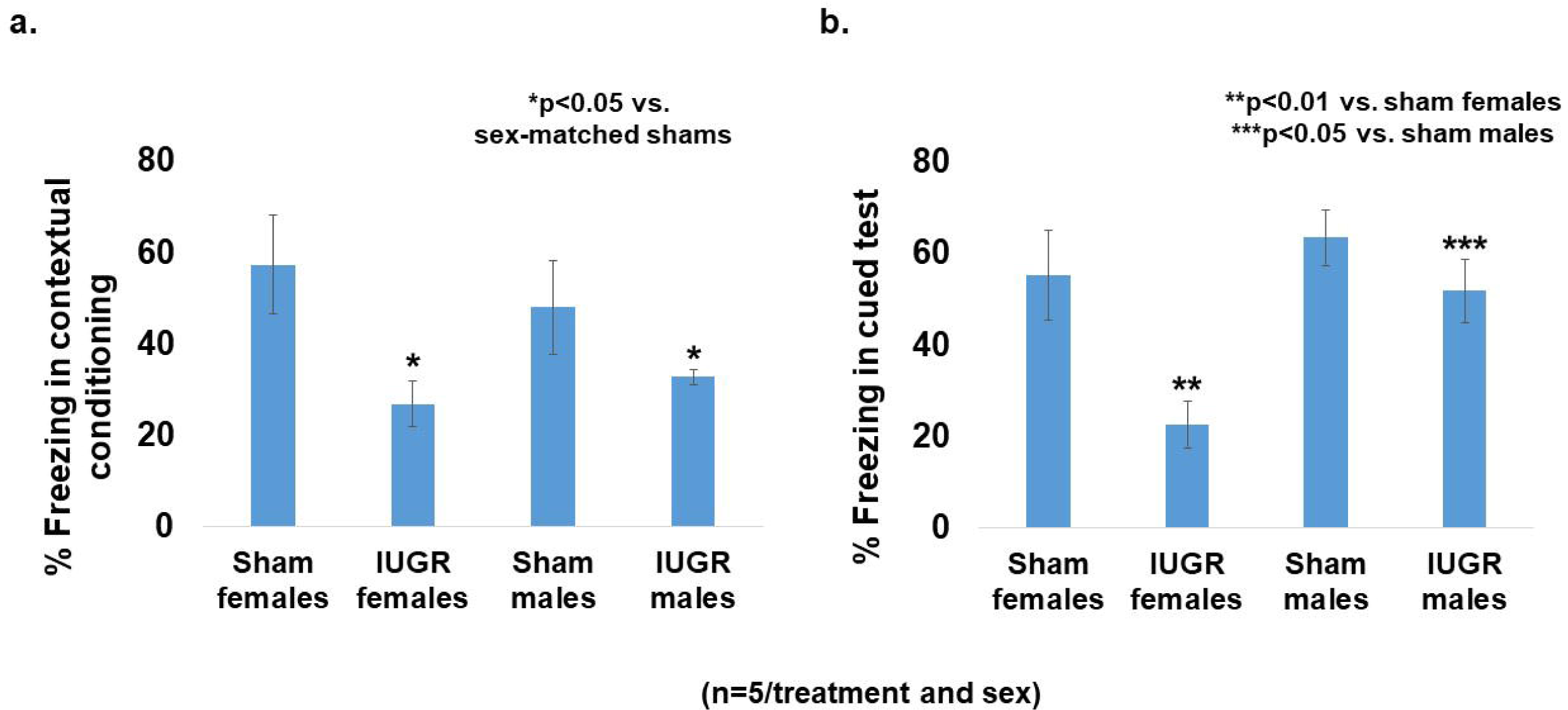
(a) Percentage of time spent freezing in contextual conditioning. IUGR offspring of both sexes spent less time freezing when the audible tone was heard but no foot shock followed. *p<0.05 compared to sex-matched sham controls. (b) Percentage of time spent freezing in cued test. IUGR offspring of both sexes spent less time freezing when placed in the same chamber in which a previous audible tone gave rise to a foot shock. **p<0.01 compared to sham females. ***p<0.05 compared to sham males. n=5 for each treatment (sham or IUGR) and sex (male or female).

#### Open field test to assess spontaneous activity and locomotion

IUGR males and females had increased total velocity of movement compared to sex-matched sham controls (7.79±0.48 cm/s in IUGR, n=16 vs. 6.57±0.41 cm/s in sham, n=18, p<0.05), which was driven by an increased velocity in the center of the field (9.19±0.71 cm/s in IUGR vs. 7.13±0.52 cm/s in sham, p<0.05) rather than in the periphery (7.07±0.43 cm/s in IUGR vs. 6.34±0.41 cm/s in sham, p>0.05). This indicates a higher speed of movements in an unprotected area. We otherwise saw no differences in the total distance traveled including distances traveled in the center or periphery, no differences in the duration of time or frequency spent in center or periphery, no difference in the latency of time moving from center to periphery, no differences in % activity in center or periphery, and no difference in rearing frequencies between sham control and IUGR mice.

#### Elevated plus maze to assess anxiety-like behavior

IUGR males and females showed increased head dip frequencies in the open arms compared to sex-matched sham controls (67.8±3.6% in IUGR, n=18 vs. 51.0±4.5% in sham, n=17, p<0.05), indicating increased exploration in unprotected areas. We otherwise saw no differences in the frequency or duration in the open vs. closed arms, no difference in the latency of time moving from open to closed arms, and no difference in stretch frequencies in open or closed arms.

#### Prepulse Inhibition (PPI) to assess pre-attentive functioning

We found a baseline sex difference with females having decreased startle amplitude compared to males (515±79 arbitrary units in females, n=19 vs. 934±122 arbitrary units in males, n=19, p<0.05). However with prepulses of 3, 6, and 12 dB above background, we found similar prepulse inhibition of acoustic startle responses between sham control and IUGR mice (data not shown), showing intact pre-attentive functioning with the paradigm tested.

#### Forced swim test to assess immobility/depressive-like behavior

Sham control and IUGR males and females showed similar frequency and duration of struggle as well as similar frequency, duration, and latency to treading water and floating.

#### Elevated bridge test to assess impulsivity and motor coordination

IUGR male and female mice showed a faster latency to cross the bridge compared to sham control mice (2.68±0.36 seconds in IUGR vs. 5.34±1.21 seconds in sham, p<0.05) but the time taken to cross the bridge was similar. These data indicate increased impulsivity and/or decreased fear.

In summary, young adult IUGR mice revealed a marked deficit in short-term implicit memory under the various testing paradigms, making a case to examine the effects of IUGR on DG neurogenesis.

### Quantification of hippocampal neurogenesis markers at E15.5

Three days after maternal hypertension, IUGR offspring of both sexes had a decreased percentage of Sox2^+^ NSPCs among the total number of DAPI^+^ cells in the ventricular zone (VZ) compared to sham controls (Figure 3a). The total absolute number of DAPI^+^ cells were similar between E15.5 sham and IUGR DG (1842±62 in sham males vs. 1754±70 in IUGR males; 1802±85 in sham females vs. 1736±61 in IUGR females, p>0.05). A 2 h EdU pulse labeling of all progenitor cells showed that IUGR offspring had a decreased percentage of proliferative cells in the VZ compared to sham controls in both sexes (Figure 3b). Surprisingly, IUGR offspring had an increased percentage of Tbr2^+^ INPs both in the marginal zone (MZ), dentate migratory stream, and dentate fimbriodentate junction (FDJ) in the E15.5 DG compared to sham controls in both sexes (Figure 3c).

**Figure 3.**
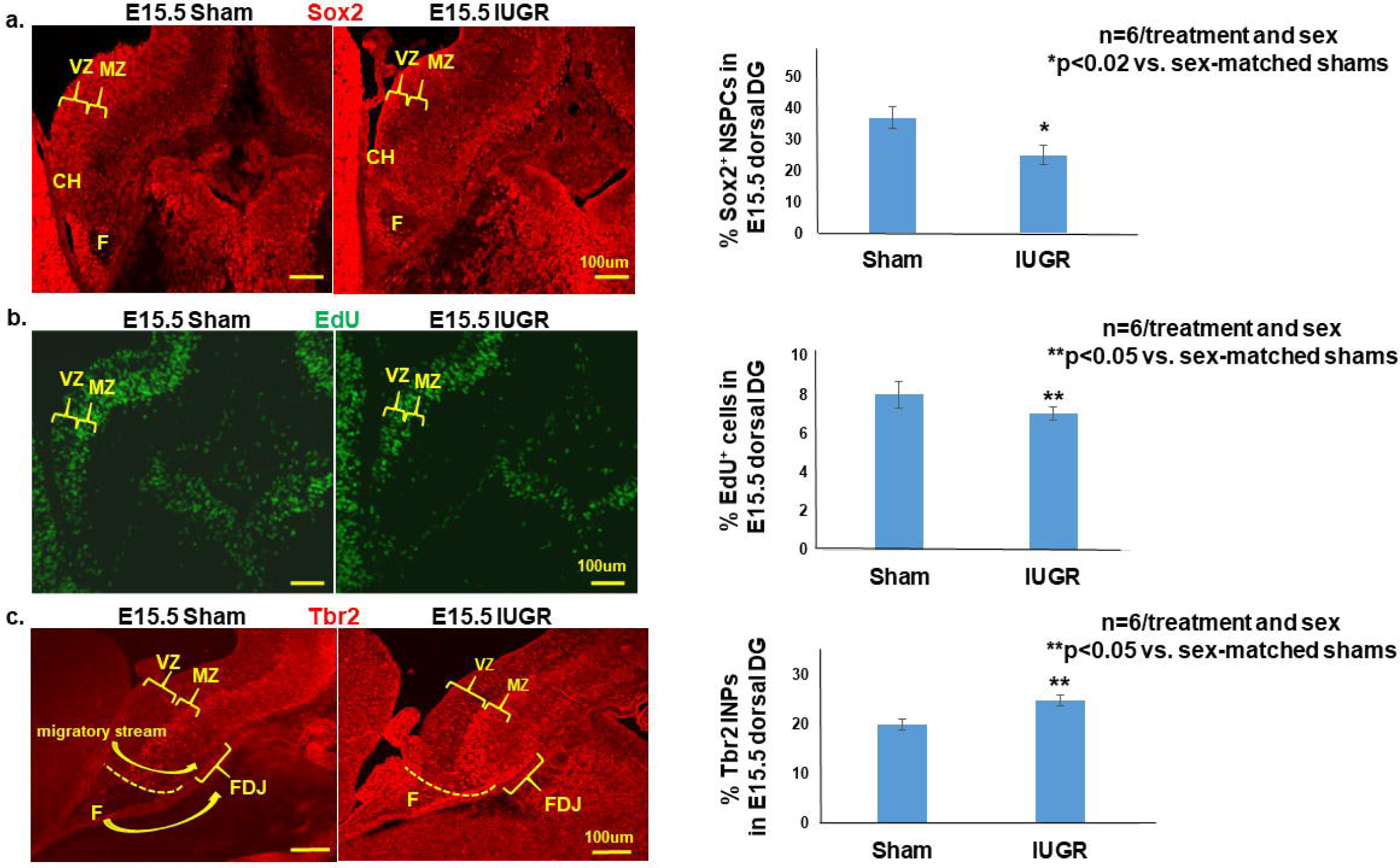
(a) Percentage of Sox2^+^ NSPCs in E15.5 dorsal DG. IUGR offspring of both sexes showed decreased percentages of Sox2^+^ NSPCs in the ventricular zone (VZ) and cortical hem (CH). (b) Percentage of EdU^+^ proliferative cells in E15.5 dorsal DG. IUGR offspring of both sexes showed decreased percentages of EdU^+^ proliferative cells, i.e. cells in S phase of cell cycle, in VZ and marginal zone (MZ). (c) Percentage of Tbr2^+^ INPs in E15.5 dorsal DG. IUGR offspring of both sexes showed increased percentages of Tbr2^+^ INPs in the MZ, migratory stream, and fimbriodentate junction (FDJ). *p<0.02 compared to sex-matched sham controls. **p<0.05 compared to sex-matched sham controls. n=6 for each treatment (sham or IUGR) and sex (male or female).

### Quantification of hippocampal DG neurogenesis markers at E19

In late maternal hypertension, IUGR offspring of both sexes continued to have a decreased percentage of Sox2^+^ NSPCs in the DG (Figure 4a, left) while proliferation was unaffected (Figure 4b, right). Important to highlight here is that the quantification of proliferative cells included the ventricular zone, dentate migratory stream, and within the dentate anlage. A difference appears to exist within the dentate anlage where IUGR had decreased EdU-labeled cells but the total percentage was overall unchanged. IUGR females at late gestation additionally showed increased percentages of Tbr2^+^ INPs, NeuroD^+^ NPs, and Prox1^+^ immature and mature granule neurons compared to sham females (Figure 4b-d). By contrast, IUGR males showed increased percentages of NeuroD^+^ NPs and Prox1^+^ granule neurons compared to sham males (Figure 4c-d). In addition to increased percentages of neuronal progenitor cells and immature/mature neurons, the localization of these cells appeared more widely distributed across the dentate plate rather than confined to the future suprapyramidal or infrapyramidal blade or the hilus.

**Figure 4.**
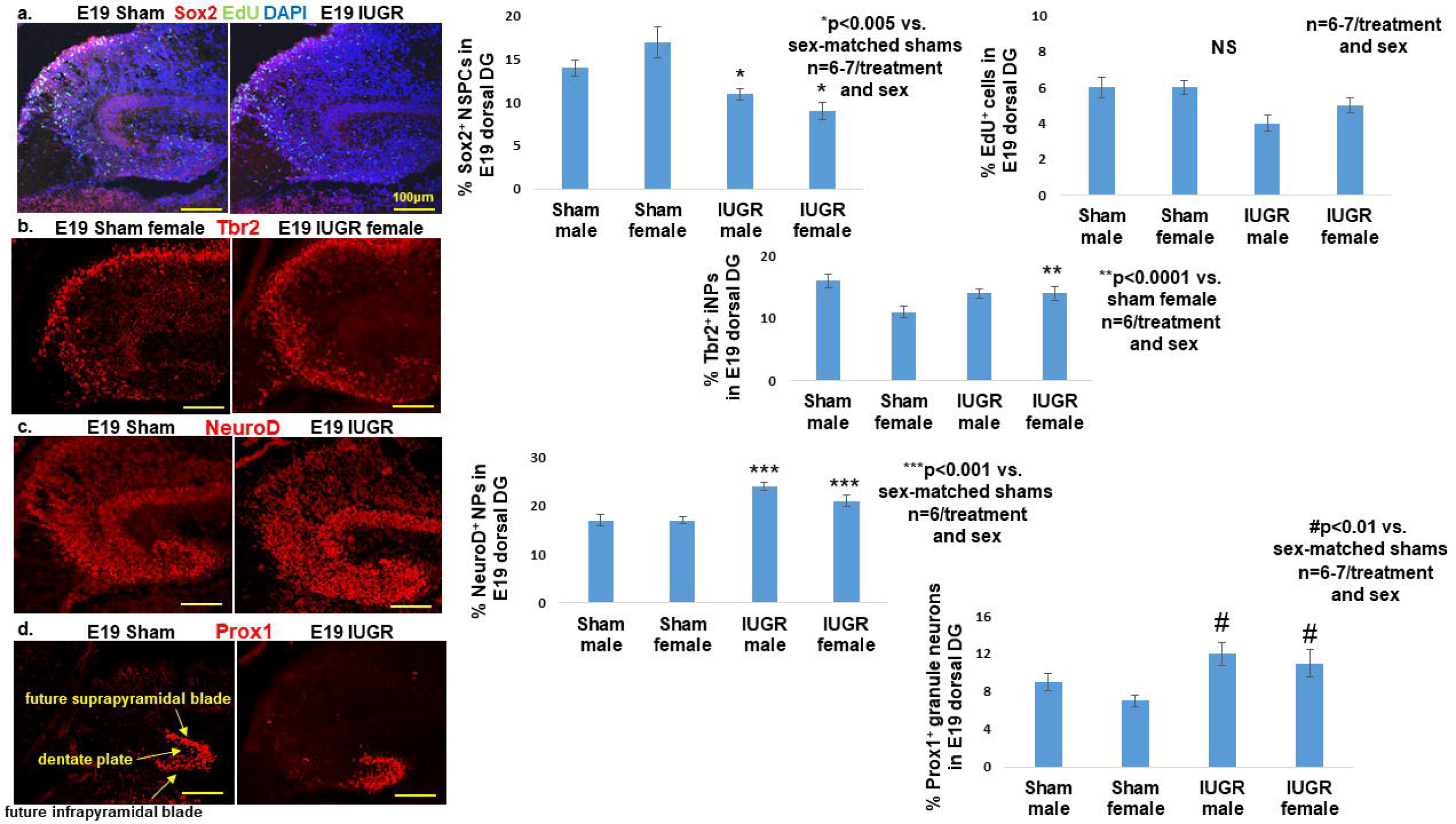
(a) Percentages of Sox2^+^ NSPCs and EdU^+^ proliferative cells in E19 dorsal DG. IUGR offspring of both sexes showed decreased percentages of Sox2^+^ NSPCs in the VZ and migratory stream. By contrast, IUGR offspring of both sexes showed similar percentages of EdU^+^ proliferative cells in VZ and migratory stream. (b) Percentage of Tbr2^+^ INPs in E19 dorsal DG. IUGR female offspring showed increased percentage of Tbr2^+^ INPs in the MZ, migratory stream, and dentate anlage. (c) Percentage of NeuroD^+^ NPs in E19 dorsal DG. IUGR offspring of both sexes had increased percentages of NeuroD^+^ NPs in dentate anlage. (d) Percentage of Prox1^+^ immature and mature granule neurons in E19 dorsal DG. IUGR offspring of both sexes had increased percentages of Prox1^+^ granule neurons in dentate plate and granule cell layers of the future supra-and infra-pyramidal blades. *p<0.005 compared to sham controls. **p<0.0001 compared to sham females. ***0<0.001 compared to sex-matched shams. ##p<0.01 compared to sex-matched shams. n=6-7 for each treatment (sham or IUGR) and sex (male or female).

### RNA sequencing of E15.5 hippocampi

Noting that IUGR promotes an imbalance towards neuronal differentiation over NSPC self-renewal, we performed RNA sequencing of E15.5 sham control and IUGR hippocampi to determine which downstream genes may be responsible for the cellular phenotype. Of the 16,873 mouse genes analyzed, 611 protein-coding gene transcripts were differentially expressed (Figure 5a). Approximately 70% of affected genes were downregulated as a result of IUGR. Ingenuity Pathway Analysis identified the canonical (Wnt/ß-catenin) and non-canonical (PCP) Wnt signaling pathways as significantly downregulated in E15.5 IUGR hippocampi (blue bars, Figure 5b). The neuropathic pain signaling pathway which has the same genes as glutamate receptor signaling plus cAMP response dependent binding protein (CREB) signaling were predominantly upregulated (orange bars, Figure 5b). Table 1 lists the protein-coding genes identified in these dysregulated pathways.

**Table 1.** List of differentially expressed protein-coding genes between E15.5 sham and IUGR hippocampi

| Gene Categories | Gene Names | Fold Changes |
| --- | --- | --- |
| Wnt and PCP signaling | *Wnt2b* | ↓3.8x |
|  | *Wnt3a* | ↓6x |
|  | *Wnt8b* | ↓2.7x |
|  | *Wnt9a* | ↓2.8x |
|  | *Wnt10a* | ↓3.3x |
|  | *Dkk1* | ↓5x |
|  | *Wif1* | ↓3.6x |
|  | *Rspo1, 2, 3* | ↓3.3x, 5.2x, 2.3x |
|  | *Sp5* | ↓2.1x |
| Glutamate receptor signaling<br>via CREB signaling | *Grm1, 5, 8* | ↑2.1x, 2.6x, 2.3x |
|  | *Grin3a* | ↑2.2x |
|  | *Gng4* | ↑3.4x |
|  | *Plcd3* | ↓2.7x |
|  | *Pla2g16* | ↓2.6x |
|  | *Abhd3* | ↓8.5x |

**Figure 5.**
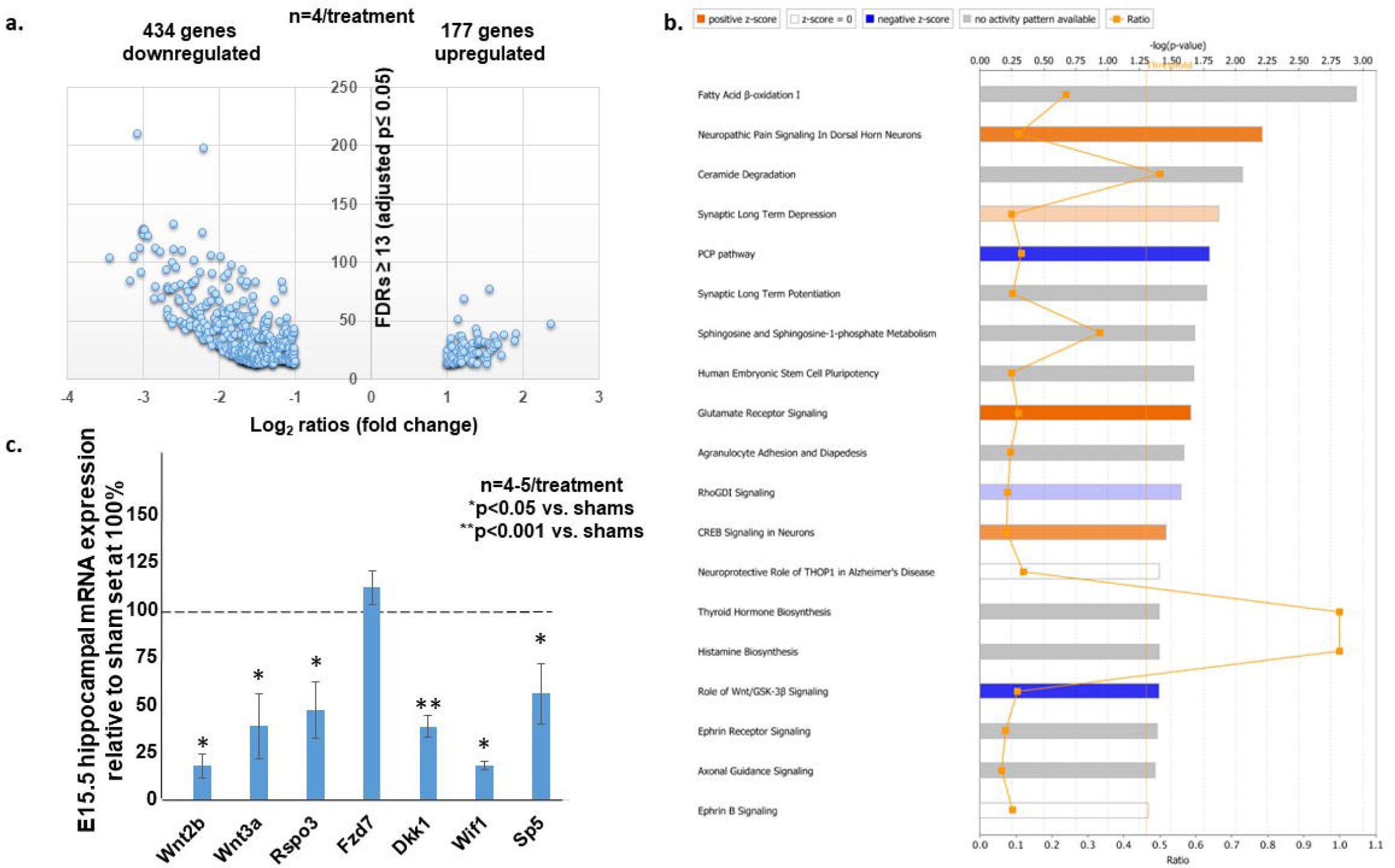
(a) Volcano plot of the 611 differentially expressed protein-coding gene transcripts in E15.5 IUGR hippocampus via RNA-seq. 434 genes (∼70%) were downregulated. The x-axis denotes log_2_ ratios which are fold changes in gene expression (log2 ratios ≥+1 or ≤-1 = two-fold increase or decrease). The y-axis denotes false discovery rates (FDRs) ≥ 13 = adjusted p value ≤ 0.05. n=4/treatment. (b) Ingenuity pathway analysis of the 611 differentially expressed genes to determine canonical pathways that were aberrantly expressed in IUGR at E15.5. Genes relating to the canonical Wnt/GSK-3β and non-canonical (PCP) pathways were significantly decreased in IUGR (blue bars denoting negative z-scores). Genes relating to the neuropathic pain signaling which list the same genes as those in glutamate receptor signaling plus CREB signaling were significantly increased in IUGR (orange bars denoting positive z-scores). Gray bars denote genes with no identifiable activity pattern. White bars denote genes in that pathway with a zero z-score. Pathways with –log(p-value) above the orange threshold line denote significance of p≤0.05. Ratios signify the number of genes detected in RNA-seq in that pathway divided by the number of genes known in that pathway. (c) Percentages of change in canonical and non-canonical Wnt (PCP) signaling pathways discovered from RNA-seq were validated with real time qPCR in E15.5 hippocampi. IUGR reduced hippocampal gene expression of Wnt2b and Wnt3a ligands, Rspo3 ligand for the PCP pathway, Dkk1 and Wif1 inhibitors, and Sp5, a readout of the canonical pathway. Sham values were normalized to 100% and IUGR-induced changes were expressed as percentages of sham values. *p<0.05 compared to sham controls. **p<0.001 compared to sham controls. n=4-5/treatment.

### Real-time qPCR of E15.5 hippocampi

Knowing the crucial role Wnt signaling plays in maintaining stem and progenitor cell identity (14) and given the predominant downregulation of this pathway in IUGR, we validated the mRNA expression of these canonical and non-canonical Wnt pathway genes in a separate cohort of E15.5 sham control and IUGR hippocampi. Similar to RNA-seq, we found that the *Wnt* ligands, *Wnt 2b* and *Wnt3a*, were decreased (Figure 5c). Similar decreases were detected in *R-Spondin 3* (*Rspo3*), a ligand for LGR4-6 receptors that activates the Wnt/PCP pathway, *Dkk1* and *Wif1* which are inhibitors of the Wnt pathway, and the *Sp5* transcription factor often used a readout of the Wnt pathway. *Fdz7* receptor, which binds Wnt ligands, in contrast, was unaltered in both RNA-seq and qPCR.

## Discussion

Children born with IUGR are at increased risk for cognitive impairment among other negative health outcomes (15-17). One area of impairment is in learning and memory, which is mediated in part by the hippocampus. Despite this known risk, no human or animal studies exist to describe the *in utero* hippocampal changes that may lead to learning and memory deficits in postnatal life. Such a lack of knowledge not only impedes our ability to counsel families about mechanisms of disease, but also hinders our ability to design therapy targeted at improving learning and memory function. Recognizing this gap, our laboratory developed a mouse model of IUGR that closely mimics human IUGR (11, 12, 18-20) in order to dissect the functional, cellular, and molecular phenotypes of the developing IUGR hippocampus.

Behavioral testing of young adult IUGR mice revealed a marked deficit in short-term implicit memory that was represented by a lack of object recognition, contextual memory, and cued memory during short-term recall. Our finding of implicit memory deficit echoes what is known in IUGR children who have been described to have poorer memory performance in multiple studies (3, 21-23). Because learning and memory deficits can be affected by other domains of behavior (24), we performed several other complementary tests that may impact hippocampal-based tasks. We detected no gross deficiency in spontaneous activity or locomotion, no signs of generalized anxiety, and intact sensorimotor gating between sham control and IUGR young adult mice. However, we have uncovered subtle changes in movement speed and impulsivity in this model that are likely attributed to other brain regions in addition to the hippocampus. The impact of IUGR on other brain regions is beyond the scope of this study but warrants future investigation.

With the demonstration of implicit learning and memory deficits, we next pursued a comprehensive characterization of embryonic DG neurogenesis. We focused on the DG because it has a distinct NSPC pool known to be regulated by environmental signals throughout life. DG neurons are the primary afferent input from the entorhinal cortex into the hippocampal formation. Furthermore, aberrant DG neurogenesis is strongly associated with learning and memory deficits (25-28). Correlating the timing of maternal hypertension to the timing of DG neurogenesis, we found that IUGR decreased the percentage of Sox2^+^ NSPCs in the ventricular zone of the cortical hem along with decreased proliferation. Contrary to our expectation, IUGR increased the percentage of Tbr2^+^ INPs that populated the marginal zone and migratory stream and in the subpial stream towards the fimbriodentate junction. Collectively, these findings suggest that Sox2^+^ NSPCs are likely precociously exiting cell cycle and becoming Tbr2^+^ INPs as a result of IUGR. As maternal hypertension continued, Sox2^+^ NSPCs in the IUGR DG continued to be decreased in percentage but retained proliferative capacity similar to sham NSPCs. IUGR males showed a slightly faster maturation into NeuroD^+^ NPs and Prox1^+^ granule cells, whereas IUGR females displayed the entire spectrum of neurogenesis markers. Our finding of accelerated neurogenesis was surprising given that postnatal studies in other IUGR animal models have shown decreased neuron number (8, 29, 30). We do not currently know whether these prematurely generated neurons survive and integrate properly into circuitry or instead undergo apoptosis. As a potential consequence of accelerated neurogenesis, embryonic Sox2^+^ NSPC depletion through symmetric differentiating divisions could pose an additional problem for the postnatal growth of the developing brain. This possibility is supported by a recent study in which Tbr2^+^ cells exhibited premature differentiation in human cortical organoids exposed to <1% oxygen, and modulators of unfolded protein response pathways could rescue this phenotype (31).

Finally, in search of the molecular pathways responsible for accelerated neurogenesis and embryonic NSPC depletion, we have identified that the canonical and non-canonical Wnt signaling pathways as potential candidates. Both pathways require Wnt ligand binding to one of the cognate receptor types, such as the frizzled genes, but after this the two pathways diverge substantially. Our RNA-seq data identified many *Wnt* ligands as decreased in IUGR. Of these, *Wnt3a* is of notable significance not only because of its greatest decrease in expression, but also *Wnt3a* expression marks the cortical hem beginning at E9.75 in mice from which induction and/or patterning of the hippocampus originates (32, 33). In *Wnt-3a* mutants, medial hippocampal fields are absent and lateral hippocampal fields are severely reduced owing to the lack of proliferative expansion of caudomedial stem and progenitor cells. In the case of IUGR, *Wnt3a* reduction could lead to Sox2^+^ NSPC reduction owing to a lack of proliferation which would be consistent with our findings. Additional support for the relevance of Wnt signaling in IUGR, we also found decreases in *Dkk1* and *Wif1* which are inhibitors of the Wnt pathway.

These Wnt inhibitors are downregulated either as a result of a negative feedback loop from the overall decreased Wnt signaling evident by decreased *Sp5* or to allow for remaining Wnt ligands to bind to maintain residual Wnt activity. In the setting of decreased NSPC proliferation, cell cycle exit would lead the IUGR brain to differentiate towards the neuronal lineage. This is evident by a significant increase in Tbr2^+^ INPs. Hodge et al. demonstrated that Tbr2 is critically required for DG neurogenesis in both developing and adult mice (34). In the absence of Tbr2, INPs are depleted despite augmented NSPC proliferation. Of particular importance, they also found that Tbr2 is enriched at T-box binding sites in the Sox2 locus to suppress Sox2 expression, suggesting that Tbr2 may promote progression from multipotent NSPC to neuronal-specified INPs by directly regulating the Sox2 gene. Once the transition to Tbr2 is established, neuronal maturation proceeds in an unhindered fashion in IUGR. The increase in glutamatergic receptor signaling could provide a mechanism for immature neurons to survive in IUGR. In developing brain, classical neurotransmitters such as glutamate exert trophic effects before synaptogenesis (35, 36). As such, the immature non-contacted cells must express functional receptors to glutamate for neuronal survival. Our RNA-seq data showing increased glutamate receptor signaling is consistent with DG granule neuron maturation and survival after NSPC cell cycle exit.

In summary, our behavioral, cellular, and molecular data lead us to conclude that short-term implicit memory deficits in young adult IUGR offspring may be associated with aberrant DG neurogenesis via decreased Wnt signaling in prenatal life. Downregulation of Wnt signaling in face of uteroplacental insufficiency results in cell cycle exit and premature commitment to neuronal differentiation. While accelerated differentiation may be an adaptive response of the fetus to ensure survival by producing more neurons, because DG granule neuron generation continues in the first postnatal month in mice, precocious neurogenesis could result in neuronal apoptosis, improper neuronal integration into circuitry, as well as continued exhaustion of NSPCs which are needed for an expanding tissue such as the growing hippocampus. When we validate decreased Wnt signaling as the underlying basis of IUGR-induced aberrant DG neurogenesis, future therapeutic intervention aimed at augmenting Wnt signaling may provide an avenue to negate premature neuronal differentiation and NSPC depletion in IUGR.

## Acknowledgements

We thank Andrew Sung for his technical assistance in immunofluorescent staining, image acquisition, and in open field behavioral data analysis. We thank Jessica Jung for her cryosectioning of the embryonic mouse brains. We additionally thank Marco Bortolato, PhD, for his expert guidance in the choice of behavioral tests.

## Author Contributions

Camille Fung, Ashley Brown, and Richard Dorsky had substantial contributions to conception and design. Camille Fung, Ashley Brown, Matthew Wieben, Shelby Murdock, and Jill Chang participated in acquisition, analysis and interpretation of data. Maria Dizon provided resources and supervised data acquisition and analysis to behavioral studies. Camille Fung, Maria Dizon, Richard Dorsky drafted and edited the article for important intellectual content. All authors had final approval of the version to be published.

## Statement of Financial Support

M.D. is supported by NINDS R01 NS086945. Majority of funding for this work was provided by the division of Pediatric Neonatology at the University of Utah School of Medicine.

## Disclosure

All authors declare no conflict of interest.

## Notes

Disclosure statement: None of the authors have any financial ties to products in this study or potential/perceived conflicts of interest.

### Competing Interest Statement

The authors have declared no competing interest.

